# Interspecific transfer of root-exuded specialized metabolites coincides with chromatin regulation and systemic chemical defense in rice

**DOI:** 10.64898/2026.07.10.737713

**Authors:** Laura Mathieu, Rémi Pélissier, Inès Bénameur, Nicolas Poncelet, Adam Rochepeau, Claudia Rouveyrol, Pierre Pétriacq, Jean-Benoit Morel, Louis-Valentin Méteignier

## Abstract

- Benzoxazinoids are indole-derived specialized metabolites released into the soil through root exudates. Initially studied for their allelopathic and toxic effects, they are now recognized as broad regulators of plant–organism interactions, including microbiome-mediated pathogen resistance. However, whether benzoxazinoid-containing root exudates can directly influence disease susceptibility in neighboring plants remains unclear.
- Using an agriculturally relevant rice–maize co-culture system, we show that benzoxazinoids naturally exuded by maize roots are taken up by rice roots and are associated with reduced rice blast disease in leaves.
- This protection occurs without detectable benzoxazinoid accumulation in rice leaves, constitutive immune activation or decreased plant height. Instead, benzoxazinoid uptake by rice roots is associated with chromatin hyperacetylation, increased expression of key phenylpropanoid biosynthetic genes and broad metabolic reprogramming.
- These responses extend systemically to leaves, where rice establishes a defense-related chemical state distinct from the systemic acquired resistance previously observed in benzoxazinoid-dependent, microbiome-mediated plant–soil feedbacks.
- Our findings support a model in which specialized metabolites exuded by one crop species are acquired by a neighbouring species and trigger chromatin-associated metabolic reprogramming linked to systemic chemical defence. This study provides a molecular framework connecting plant–plant chemical interactions, root exudation, chromatin regulation and disease susceptibility.

## Introduction

Plants influence neighboring species through multiple above- and belowground processes, yet the molecular mechanisms underlying root-exudate-mediated interactions remain poorly understood. More broadly, although the benefits of plant diversity for agroecological and ecosystem services through belowground processes are well recognized (Cappelli *et al*., 2022), the identity of the molecular cues involved, their trafficking between neighboring plant species, and the mechanisms by which these biochemical fluxes shape plant physiology remain largely unresolved (Mathieu *et al*., 2024). This gap is particularly relevant in maize-based diversification systems, which are widely used and can improve yield and grain quality (Li *et al*., 2023). Maize is a major source of benzoxazinoids (BXs) released through root exudates, which are indole-derived specialized metabolites historically studied for their phytotoxic and defensive properties (Belz & Hurle, 2005; Rice *et al*., 2012; Wu *et al*., 2022; Knoch *et al*., 2022). Recent studies have expanded the functional repertoire of BXs beyond allelopathy and toxicity. In contrast to reductionist assays using single purified BXs at µ-to-mM concentrations, maize-conditioned soils containing natural BX cocktails at the nM scale (Stengele *et al*., 2026) can improve subsequent crop productivity (Gfeller *et al*., 2023, 2024). These effects have often been attributed to BX-mediated restructuring of soil microbial communities (Hu *et al*., 2018a; Cotton *et al*., 2019; Stengele *et al*., 2026). For instance, naturally-conditioned soils with BXs shaped a beneficial microbiome that enhanced growth, systemic acquired resistance-related gene expression, and pathogen resistance in *Arabidopsis thaliana* (Stengele *et al*., 2026). Microbes also influence BX synthesis, as a microbe-specific enzymatic activity is required to generate the full spectrum of BXs chemodiversity (Thoenen *et al*., 2024a). However, BX-containing soils can modulate plant performance without involving microbiome changes, suggesting that BXs may also directly regulate receiver plant physiology (Kong *et al*., 2018; Zhou *et al*., 2018; Gfeller *et al*., 2023). This raises the possibility that compounds historically viewed as toxic or defensive may, at natural interspecific exposure levels and/or in specific biological contexts, elicit adaptive responses in neighboring plants consistent with a hormesis-like logic (Agathokleous *et al*., 2020).

In line with this possibility, recent studies showed that BXs released by donor plants can be taken up by roots and accumulate in tissues of diverse receiver species (Hazrati *et al*., 2020; Hama *et al*., 2022, 2024, 2026). In addition, BX signaling in maize leaves regulate callose and cathecol metabolites acumulation that participate in plant defenses (Ahmad *et al*., 2011; Richter *et al*., 2024). Yet, a central question remains unanswered: can BX-containing root exudates internalized within neighbor tissues trigger systemic metabolome reprogramming and disease-related phenotypic changes in distant organs of a neighboring crop species? This question is particularly compelling because BXs influence gene regulation beyond their local biochemical or toxic activity. In *Arabidopsis thaliana*, *in vitro* treatments with single purified BXs induced genome-wide transcriptional changes associated with allelopathic growth inhibition, including upregulation of biotic and abiotic stress gene ontologies (Baerson *et al*., 2005; Venturelli *et al*., 2015). BX-specific gene expression changes were associated with histone hyperacetylation, indicating that BXs or BX-derived responses act through chromatin regulation (Venturelli *et al*., 2015). However, whether naturally exuded BX cocktails transferred between crop species can trigger chromatin-associated reprogramming in a receiver plant, and whether such reprogramming is associated to local and systemic metabolome and phenotypic changes remained unknown.

Here, we use an agriculturally relevant rice-maize co-culture system (Kombiok *et al*., 2012; Tuti *et al*., 2022; Erythrina *et al*., 2022; Shahidullah *et al*., 2024) to test whether maize-derived BXs can act as interspecific chemical cues associated with rice physiological reprogramming beyond the root interface. We show that BX-containing maize root exudates are associated with suppression of the disease caused by *Magnaporthe oryzae*, a major threat to rice production (Savary *et al*., 2019). Combining genetic, chromatin, and metabolomic approaches, we provide evidence supporting that maize BXs taken up by rice roots are associated with root chromatin and metabolism activation, and establish a systemic defense-related chemical state in leaves. These findings support a model in which specialized metabolites released by one crop species can be acquired by a neighboring species and trigger chromatin-associated metabolic reprogramming linked to decreased disease susceptibility, hence revealing how crop diversification may harness hormesis-like chemical interactions to shape plant health.

## Material and methods

### Plant growth conditions

*Oryza sativa* cv. Nipponbare (NPB) seeds were cultured in commercial soil (Neuhaus N2, 3.5 g/L Basacote 14-3-19) in pure or in species mixture with B73 (wild-type) or a homozygous near-isogenic B73 *bx1* mutant (Maag *et al*., 2016a). Maize seeds were sown one week later than rice to mitigate the different growth rate between the two species. Co-cultures of eight NPB plants in pure or four NPB with one maize plant was then allowed for two additional weeks in the same pot (8 x 8 x 8 cm) at 16h light-8h dark cycles at 23-27°C and 60% relative humidity for 6 days. Pots containing less than three NPB plants were not analyzed. The total height of rice plants (spread-out leaves) was recorded after two weeks of co-culture, on the day of infection. In a set of experiments, soil was conditioned for three weeks with eight NPB plants or a single B73 WT or mutant maize plant. The resulting soil was diluted half with unconditioned soil before growing eight NPB plants in pure for three weeks before infection.

### *Magnaporthe oryzae* infection and symptom quantification

*Magnaporthe oryzae* GUY11 strain virulent on NPB (Zhang *et al*., 2018) was grown for 10 days on rice flour agar medium (20 g/L of rice seed flour, 15 g/L of agar, 2.5 g/L of yeast extract) under fluorescent light for 12h/day at 26°C for 10 days. Conidia were harvested by flooding the plate with 5 mL of sterile distilled water. Plant trays containing 15 randomized pure and WT/mutant co-cultures were sprayed with a solution containing 100,000 conidia/mL, 0.5% gelatin and 0.1% Tween-20 at a rate of 25,000 conidia per three-week-old rice plant. Inoculated plants were kept at 24°C at 100% relative humidity for 24h in the dark before going back to growth chambers set at 16h light-8h dark cycles at 23-27°C and 60% relative humidity for 6 days.

The last fully developed infected leaf of four infected NPB plants per pot was used for disease scoring using the machine learning tool LeAFtool (https://github.com/sravel/LeAFtool) that was trained to discriminate “healthy” from “disease” pixels to calculate a % of diseased leaf area. For co-cultures and plant-soil feedbacks, 4 or 2 independently replicated experiments, each comprising 6 pots (biological replicates) per co-culture condition, were used respectively.

### RNA analysis

Rice tissues from a single pot were harvested and pooled per independent biological replicate. Tissues were harvested either at 0h before infection, or 48 hours after mock or inoculation treatments. Data from three independent biological replicates were used. Total RNAs were extracted from 100-200 mg of tissue powder with TRIzol™ and further purified with phenol:chloroform:isoamyl alcohol (25:24:1). Five µg of RNA were used for cDNA synthesis with M-MLV reverse transcriptase (Promega) with a mix of oligo dT and random primers following the manufacturer instructions. cDNA was diluted half in 1X GoTaq® qPCR Master mix with specific primers (Table S1) at 250 nM and expression data was acquired in technical triplicate with an LC480 lightcycler. Relative expression was calculated by the ΔΔCt method with *OsUBQ* as a reference gene and the pure condition as control.

### Metabolomic data acquisition

At 0h before infection, rice roots were extensively washed in 4 consecutive water baths to remove all soil material and maize root tissues, briefly dried on soaking paper and flash frozen in liquid nitrogen. Roots and leaves were ground in liquid nitrogen and freeze-dried for 48 hours. Rice tissues from a single pot were harvested and pooled per biological replicate, and 5 independent biological replicates were analyzed per co-culture condition. Ten milligrams of dried leaf and root tissue were extracted with ethanol for metabolomic profiling of semi-polar metabolites, including both primary and specialized metabolites, and the resulting extracts were analyzed by UHPLC-HRMS equipped with an electrospray ionization (ESI) source operating in negative ion mode, as previously described (Dussarrat *et al*., 2022; Pinheiro Costa Pimentel *et al*., 2026). The LC-MS injection sequence was randomized and included 36 biological samples (3 co-culture conditions × 2 tissues, n = 5; WT and mutant maize roots, n = 3), 8 extraction blanks (prepared without biological material to identify potential contaminants), and 9 quality control (QC) samples prepared by pooling 50 μL from each biological sample and standard. QC samples were used to correct for signal drift during long analytical batches and calculate the coefficient of variation for each metabolomic feature, allowing retention of only the most robust features for downstream chemometric analyses (Broadhurst *et al*., 2018). Raw LC-MS data were processed following the DIA MS2 deconvolution method using MS-DIAL software (v. 4.9; (Tsugawa *et al*., 2015)) following optimized parameters. Annotations were performed based on MS1 spectra and MS2 DDA fragmentation information using FragHUB (Dablanc *et al*., 2024) and an in-house authentic compound library for level 1 annotation. Thus, annotation of differentially accumulated metabolites resulted from MS-DIAL screening of the MS1 detected exact HR m/z and MS2 fragmentation patterns (Tsugawa *et al*., 2015). Additionally, the InChiKeys of annotated features were employed within ClassyFire to generate a structural ontology for chemical entities. Curation of 6404 raw metabolomic signals (SN > 10, CV QC < 30%) resulted in 4337 LC-MS features. Among these, 1172 remained unidentified, having no match with either MS1 or MS2.

### Data analysis and visualisation

The 3165 annotated features were further filtered (IQR=40%) and normalized (sample median, cube root transformation, Pareto scaling) for normality and comparison purposes and underwent multivariate statistical analyses using MetaboAnalyst (v6.0) (Pang *et al*., 2022). To identify metabolites responsive to benzoxazinoids while controlling for tissue-specific metabolic variation, we applied a linear modeling framework implemented in MetaboAnalyst (v6.0). Benzoxazinoid status was modeled as a categorical factor with two levels: non-exposed (rice plants grown in pure culture or co-cultured with *bx1*, which produces residual levels of benzoxazinoids) and exposed (rice plants co-cultured with B73). Tissue identity was included as a covariate to account for intrinsic metabolic differences between sample types and thereby isolate benzoxazinoid-associated effects independently of tissue context. For each metabolite, the following model was fitted:

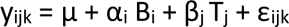

where y_ijk_ is the metabolite intensity, μ represents the mean metabolite intensity in the non-exposed reference group, α_i_ estimates the effect of benzoxazinoid exposure, β_j_ corresponds to the effect of tissue, and εᵢ_jk_ denotes residual error assumed to be normally distributed. The primary contrast tested corresponded to benzoxazinoid-exposed versus non-exposed plants, adjusted for tissue identity. Statistical significance was determined using raw P-values with a threshold of P ≤ 0.05.

Boxplots were generated in Python using Matplotlib. Boxes represent the interquartile range (IQR), with the central horizontal line indicating the median. Whiskers extend to the most extreme observations within 1.5 × IQR of the lower and upper quartiles. Individual biological replicates are shown as black points, with a small horizontal jitter applied to reduce overlap. The arithmetic mean is indicated by a hollow magenta diamond.

Non-parametric Mann-Whitney U test followed by Benjamini-Hochberg correction was applied to compare ≤ 2 conditions. To compare ≥ 3 conditions, ANOVA and Tukey HSD correction was applied.

### Chromatin analysis by western blot and ChIP-qPCR

Rice tissues from a single pot were harvested and pooled per biological replicate. Western blot, nuclei and chromatin immunoprecipitation-qPCR were performed essentially as in (Méteignier *et al*., 2022) with the following changes in the crosslinking and chromatin fragmentation steps. The crosslinking step was performed on 1.25g ground tissues resuspended in 25mL of 60 mM HEPES pH8, 1 M sucrose, 5 mM KCl, 5 mM MgCl2, 5 mM EDTA, 0.6% Triton X-100, 0.4 mM PMSF, 1X complete mini EDTA-free protease inhibitor from Roche, 1% formaldehyde, for 10 minutes at room temperature. Crosslinking was quenched by adding 5 mL of 2 M glycine and further incubation 5 minutes at room temperature. Sonication of nuclei preparation was achieved with a probe sonicator (Vibra cell 72434, Bioblock Scientific) set at 50% power, 5s ON/5s OFF cycles repeated 24 times to reach a fragmentation pattern ranging from 300 to 600 bp. Anti-H3 and -H3K27ac from Abcam (ab1791 and ab4729, respectively) and HRP-conjugated goat anti-rabbit secondary antibodies from Millipore (12-348) were used.

## Results

### Benzoxazinoid-containing maize root exudates decrease rice blast severity without obvious changes in growth or defense gene expression

We first assessed whether maize could modulate rice blast disease caused by the fungus *Magnaporthe oryzae* in rice co-cultured for two weeks. Rice plants (Nipponbare cultivar-NPB) grown with B73 maize exhibited a significant ∼4-fold decrease in disease symptoms compared to rice grown in pure conditions after two weeks of co-culture (Figure 1).

**Figure 1.**
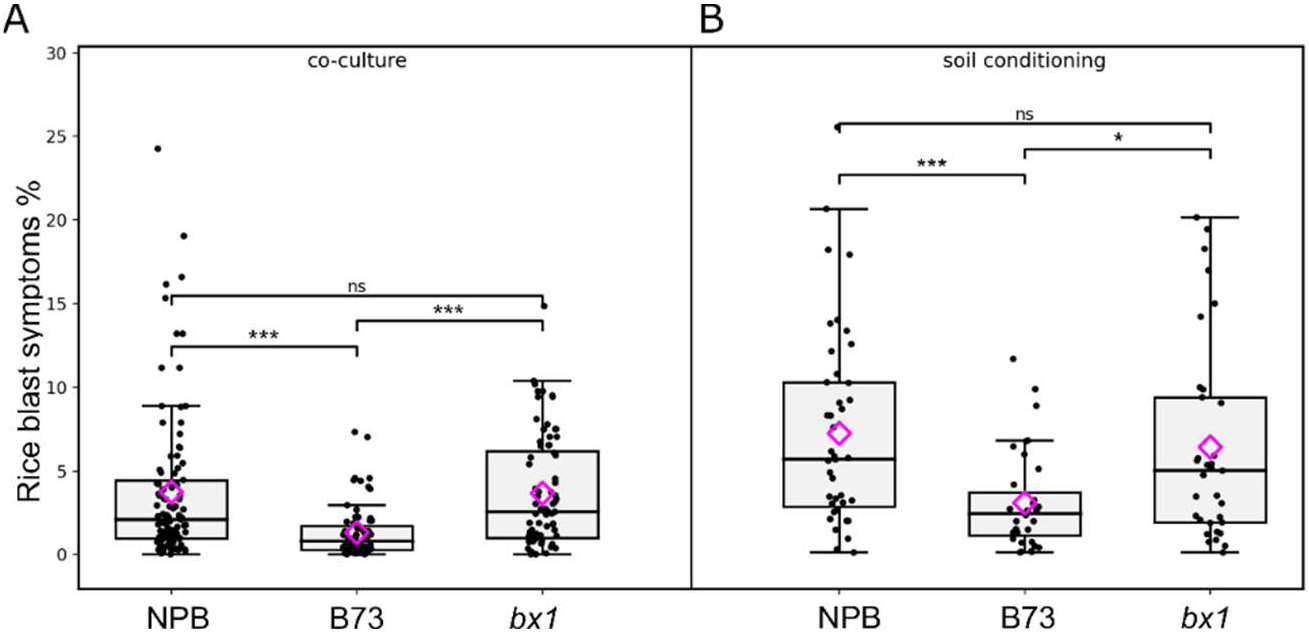
Effect of BX-containing maize root exudates on rice blast disease. **A.** *Oryza sativa* cv. Nipponbare (NPB) grown in pure or co-cultured with the B73 wild-type maize or the B73 *bx1* mutant were infected with the *Magnaporthe oryzae* GUY11 strain. Disease symptoms were quantified 6 days post-infection and normalized against the surface of leaf without symptoms. Boxplots represent values from 4 independent experiments. Magenta diamond indicates the mean. * P<0.05, *** P<0.001 according to pairwise Mann-Whitney U test with Benjamini-Hochberg correction. **B.** Conditioned soil with NBP, B73 or *bx1* for three weeks was diluted half with unconditioned and was used to grow NPB in pure. Disease infection and scoring were performed as in (A). Boxplots represent values from 2 independent experiments. Magenta diamond indicates the mean. * P<0.05, *** P<0.001 according to pairwise Mann-Whitney U test with Benjamini-Hochberg correction.

To test whether maize BXs are involved in this protective effect, we used the *bx1* maize mutant, which is strongly decreased in BX accumulation and exudation (Maag *et al*., 2016b). Unlike B73, co-cultivation with the *bx1* mutant did not reduce rice blast symptoms compared to pure conditions (Figure 1A). These results indicate that maize BXs contribute to the reduced susceptibility of rice to *M. oryzae*. To confirm the role of BXs contained in root exudates in this disease-suppressive effect, we conditioned soil either with NPB, B73, or B73 *bx1* for three weeks, and subsequently grown NPB in pure on each soil condition. We confirmed the BX-dependent suppressive effect on rice blast disease, albeit to a lesser extent (Figure 1B). These results support a role for BX-containing maize root exudates in suppressing the severity of rice blast disease in leaves.

We then examined whether BX-induced disease suppression was at the expense of a trade-off on growth, or was associated with constitutive or enhanced activation of rice defenses. Co-cultivation with maize (either B73 or *bx1*) did not affect total rice plant height (Figure S1A). At 0h before infection, gene expression analyses revealed modest transcript changes: *DEP1* (growth marker), *EXT-like* (induced systemic resistance marker), and *JAZ1* (jasmonate signaling) were slightly increased, *PR10a* (immunity marker, SA-responsive) was decreased, and *PAL1* (immunity marker, SA-responsive) and *CeBIP* (immunity marker, chitin-responsive) remained unchanged in rice leaves grown with B73 maize compared to pure conditions (Figure S1B). Forty-eight hours upon pathogen inoculation, all tested defense-related genes were similarly induced across conditions (pure culture, B73 co-culture, and *bx1* co-culture), with comparable basal expression levels in mock-treated rice (Figure S1C). We conclude that BXs do not strongly affect growth and defense gene expression before and after infection, supporting that alternative mechanisms underly BX-mediated disease suppression.

### Benzoxazinoids are transferred from maize to rice roots during co-cultivation

We next asked whether BXs can be readily detected in rice tissues. Using fragmentation patterns, we successfully matched nine experimental metabolite features against BX references considering the whole dataset including B73 maize roots (Figure S2A). As expected, heatmap clustering of samples based on BX relative levels confirmed high accumulation in roots of B73 maize, whereas BX levels were much lower but not absent in *bx1* roots (Figures S2B). Interestingly, root samples of rice co-cultured with B73 clustered closer to *bx1* root samples than to other samples (Figures S2B).

Analysis of BX accumulation in rice tissues revealed that these nine distinct BXs, for which the biosynthetic pathway is absent in rice (Dick *et al*., 2012; Wu *et al*., 2022) and thus not detected in pure condition, were readily detected in the roots of rice plants co-cultivated with B73 (Figure 2). Relative quantification showed that 20-30% of BX levels present in wild-type maize roots (with an average relative accumulation of ∼5 for all detected BXs in BX73 maize) were found in rice roots when co-cultivated with B73 maize (with an average relative accumulation of ∼2 for all detected BXs in rice) (Figure 2). In contrast, we did not detect any of the confirmed BXs in rice leaves (Figure 2). We conclude that a BX cocktail is transferred from maize to rice roots, supporting the hypothesis of a direct chemical interaction between the two species in the rhizosphere, and arguing against direct toxicity of the cocktail on *M. oryzae* as evidenced by their absence in leaves.

**Figure 2.**
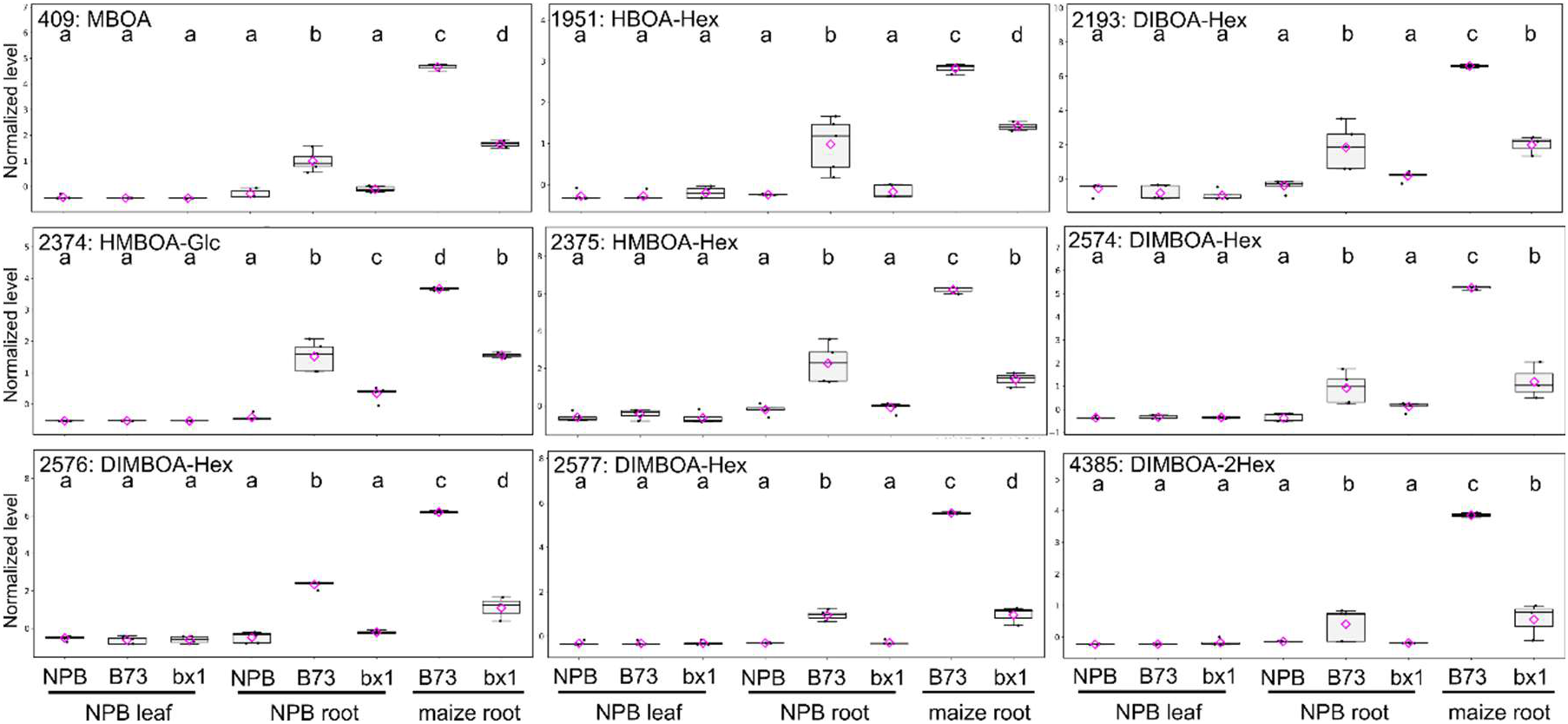
Relative quantification of benzoxazinoids in leaves and roots of rice in co-culture. Accumulation in maize roots (B73 and *bx1*) grown in pure culture is also shown. On the day of infection, rice roots from plants grown with NPB, B73 or *bx1* were thoroughly washed in water, dried on soaking paper, and freeze-dried before HPLC-MS/MS analysis. Benzoxazinoids confirmed by caomparing fragmentation spectra with references (Figure S2A) are shown here. Magenta diamond indicates the mean. Different letters indicate statistical differences based on ANOVA followed by Tukey HSD post-hoc test.

### Benzoxazinoid-dependent metabolic augmentation and contrasted effects on specialized metabolism

To explore the molecular responses of rice to the uptake of BX-containing maize root exudates, we performed untargeted metabolomic analysis of roots and leaves of rice grown in pure culture or in co-culture with B73 or *bx1* prior to inoculation. Heatmap visualisation of the 4337 high-quality metabolite features indicated clear clustering of roots and leaves and good reproducibility between independent biological replicates per condition (Figure S3). Linear modeling of BX effects on metabolite quantity with tissue effects as a fixed factor revealed 518 BX-responsive metabolites (P<0.05, Table S2), among which 60 were significant regardless of tissue correction and 458 were significant only after correcting for tissue-wise effects (Figure 3A), indicating important tissue-specific effects. We first inspected the level of jasmonic acid and catechol, known markers of BX signaling (Hu *et al*., 2018b; Richter *et al*., 2024). Jasmonic acid and catechol were significantly induced in a BX cocktail-dependent manner, particularly in roots (Figure S4A-B). Among the 518 differentially accumulated metabolites, core metabolites such as fatty acyls (Figure S4A) and carboxylic acids (Figure S4C) were highly enriched (Figure 3B) along with a number of specialized metabolic pathways linked to phenolic compounds such as benzenes (Figure 3B and S4B).

**Figure 3.**
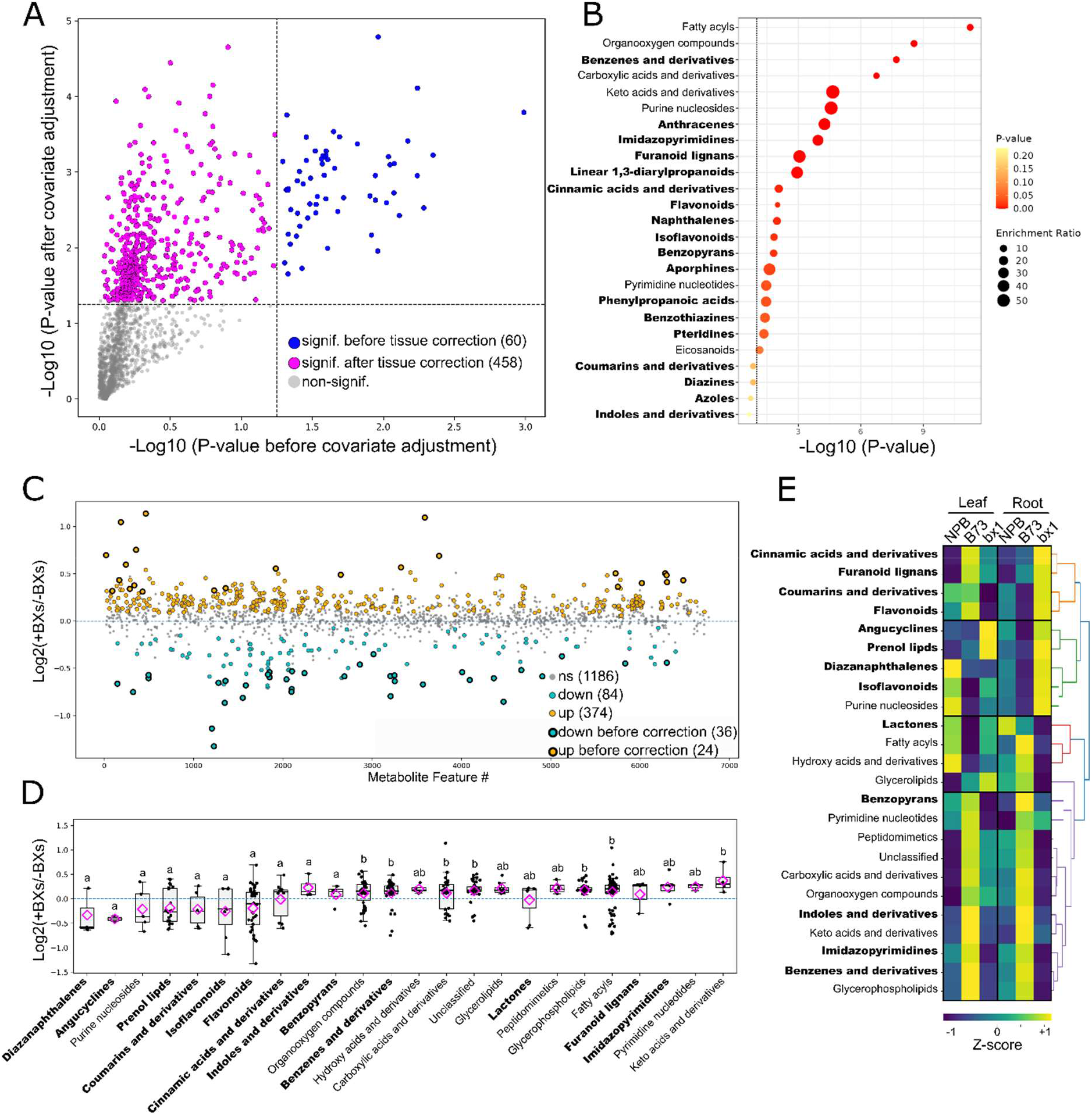
Analysis of benzoxazinoid-dependent metabolome reprogramming in rice. **A.** A generalized linear model with tissue correction as a fixed effect identified 518 rice metabolite features sensitive to the presence of benzoxazinoids. A vast majority were significant after tissue correction, highlighting tissue-specific changes. **B.** Metabolite structural class enrichment test using ClassyFire structural database for the 518 metabolite features identified by linear modelling. **C.** Manhattan plot of metabolite log2 fold changes in response to benzoxazinoids for metabolite features with assigned compound names. **D.** Structural classes possessing at least 2 features with a significant log2 FC ≥ |0.2| in response to benzoxazinoids were analyzed. Structural classes in bold indicate specialized metabolites. Magenta diamond indicates the mean. Different letters indicate statistical differences according to ANOVA followed by Tukey HSD post-hoc test. **E.** Average tissue-wise Z scores were calculated for each structural class shown in D and were clustered by Euclidean distance and average linkage according to their accumulation pattern across tissues and co-culture conditions.

Strikingly, most of the BX-responsive metabolites were upregulated (374 up *versus* 84 down, Figure 3C). Indeed, structural ontologies enriched in core metabolism were significantly more induced by BXs in comparison to most specialized metabolism pathways, except for benzene phenolic compounds (Figure 3D). In line with that, upregulated pathways in the metabolic network were enriched for core metabolism (Figure S5A), while the downregulated pathways were significantly enriched for flavonoid biosynthesis (Figure S5B). This pattern further argues against a general growth defect, which would be expected to coincide with a broad suppression of core metabolism.

To explore tissue-specific regulations, we further analyzed the average activity of each metabolic pathway in roots and leaves separately, by averaging the tissue-wise Z score of each of the 518 BX-respponsive metabolite feature per structural ontologies (Figure 3E). Structural ontologies segregated into 4 clusters. As expected, the largest cluster (dendrogram in purple) corresponded to pathways equally activated by BXs in roots and leaves, including benzenes, indoles, and most core metabolism pathways (Figure 3E). In contrast, two other clusters (orange and red dendrograms) showed distinct regulation in roots compared to leaves. Notably, most specialized pathways slightly repressed or unchanged in response to BXs in roots were instead activated in leaves, including cinnamic acids and derivatives, furanoid lignans, coumarins, and flavonoids (Figure 3E). The reverse was also true for lactones, fatty acyls and hydroxy acids, mostly over-accumulated in roots, but markedly repressed in leaves. Inspecting the level of individual metabolites within each of these contrasted clusters was in line with the global activity of each pathway (Figure S6). Taken together, these results show that the BX cocktail drives a global metabolic augmentation in neighboring rice plant roots, with an intriguing contrasted effect on specialized metabolism in leaves compared to roots.

### The contrasted effect of BXs in roots and leaves is specific to specialized metabolism

To further explore tissue-specific changes in specialized metabolite accumulation, we focused on the most confidently annotated specialized metabolites in each tissue separately. Correlation analysis revealed a highly significant negative correlation between metabolite accumulation in roots of rice co-cultivated with B73 versus *bx1* maize (Figure 4A). Consistent with this, heatmap visualisation and batch metabolite levels analysis confirmed the global increase in root metabolites in the presence of B73 in comparison to pure, which was lost in co-culture with *bx1* maize (Figures S7A-B).

**Figure 4.**
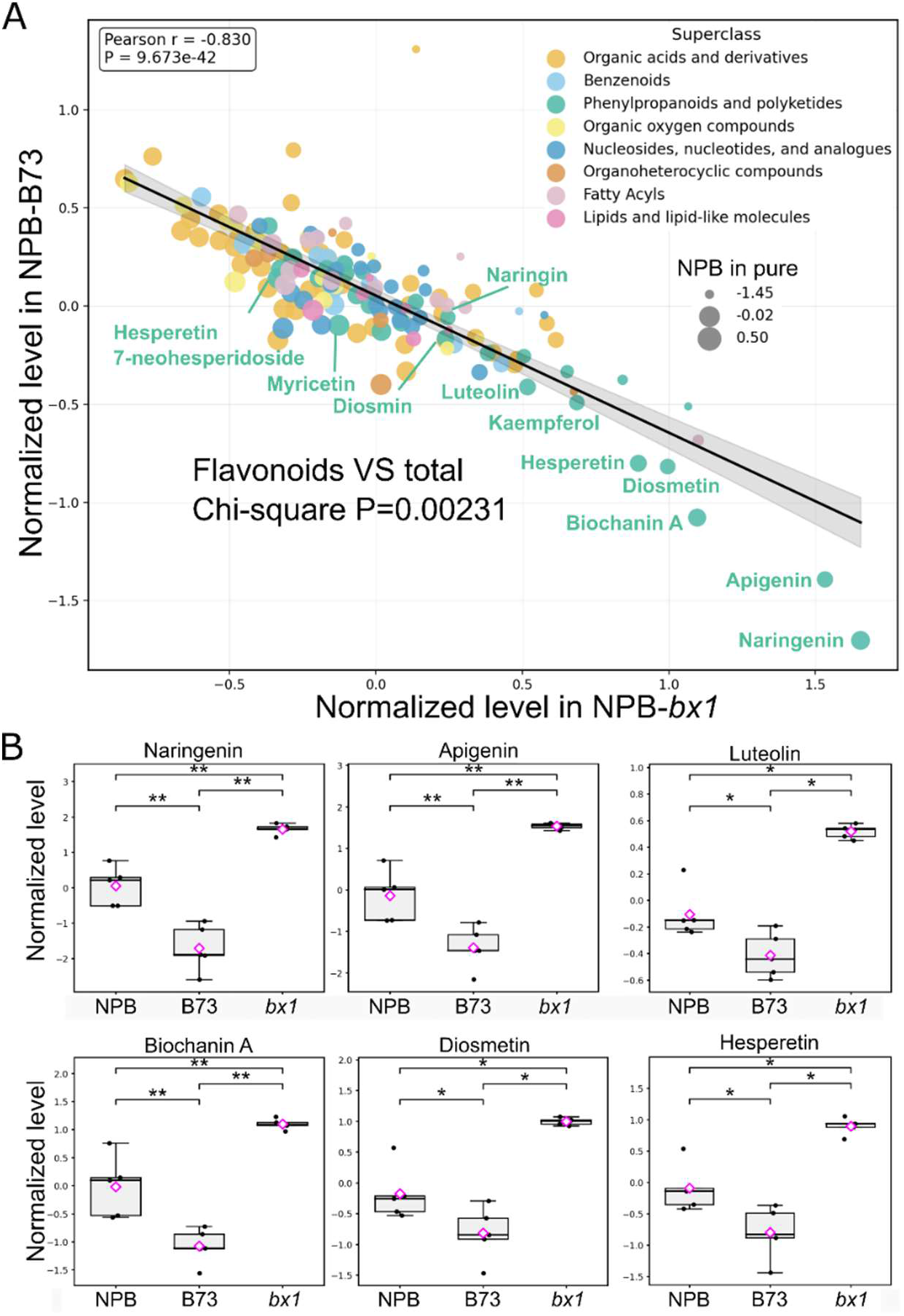
Metabolomic responses of rice roots to a BX cocktail. **A.** Normalized metabolite intensities in Oryza sativa cv. Nipponbare (NPB) roots co-cultured with wild-type maize (B73) or the benzoxazinoid-deficient mutant *bx1*. The black line represents the linear regression, and the grey shadow indicates the 95% confidence interval. Metabolites detected by HPLC-MS/MS were annotated using an authentic standard library. Each point is colored according to its ClassyFire structural superclass, and dot size reflects metabolite abundance in NPB roots grown in pure culture. Contrasted flavonoid accumulation pattern was tested against the accumulation pattern of all other metabolites by Chi-square test. **B.** Normalized intensities of representative flavonoids in rice roots. Magenta diamond indicates the mean. * P<0.05, ** P<0.01, *** P<0.001 according to Mann-Whitney U test followed by Benjamini-Hochberg correction.

In contrast with the global metabolic activation, flavonoids were strongly decreased in rice roots exposed to BXs. Inspection of individual flavonoid metabolites confirmed the strong and significant decrease of several major flavonoids in rice roots when co-cultured with B73 in comparison to the pure culture (Figure 4B). Importantly, this pattern was no longer visible in *bx1* co-cultures, indicating that BXs, although activating global metabolism, specifically decrease tissular flavonoid contents in rice roots.

Similar analysis revealed a distinct response to BX perception in leaves than in roots. A weak negative correlation was still observed between metabolite profiles of rice grown with B73 versus *bx1* maize (Figure 5A). Indeed, similar to what was observed in roots, global metabolic activity was significantly increased in rice leaves in response to B73 co-culture (Figures S8A-B). However, in contrast to roots, phenylpropanoids were enriched in over-accumulated metabolites (Figure 5B). Notably, the specific repression of flavonoids was no longer observed (Figure 5A). Instead, flavonoids such as Myricetin and Diosmin and other more simple phenolics ’(*e.g.* gallic acid,4-hydroxy-3-methylbenzoic acid) were increased in response to BXs (Figure 5C). Taken together, these results indicate that BXs induce a strong positive shift in the root metabolome, with the exception of flavonoids, while leaves undergo more complex metabolic regulations towards accumulation of defense specialized metabolites including flavonoids and simple phenolics (Figure 3E and Figure 5).

**Figure 5.**
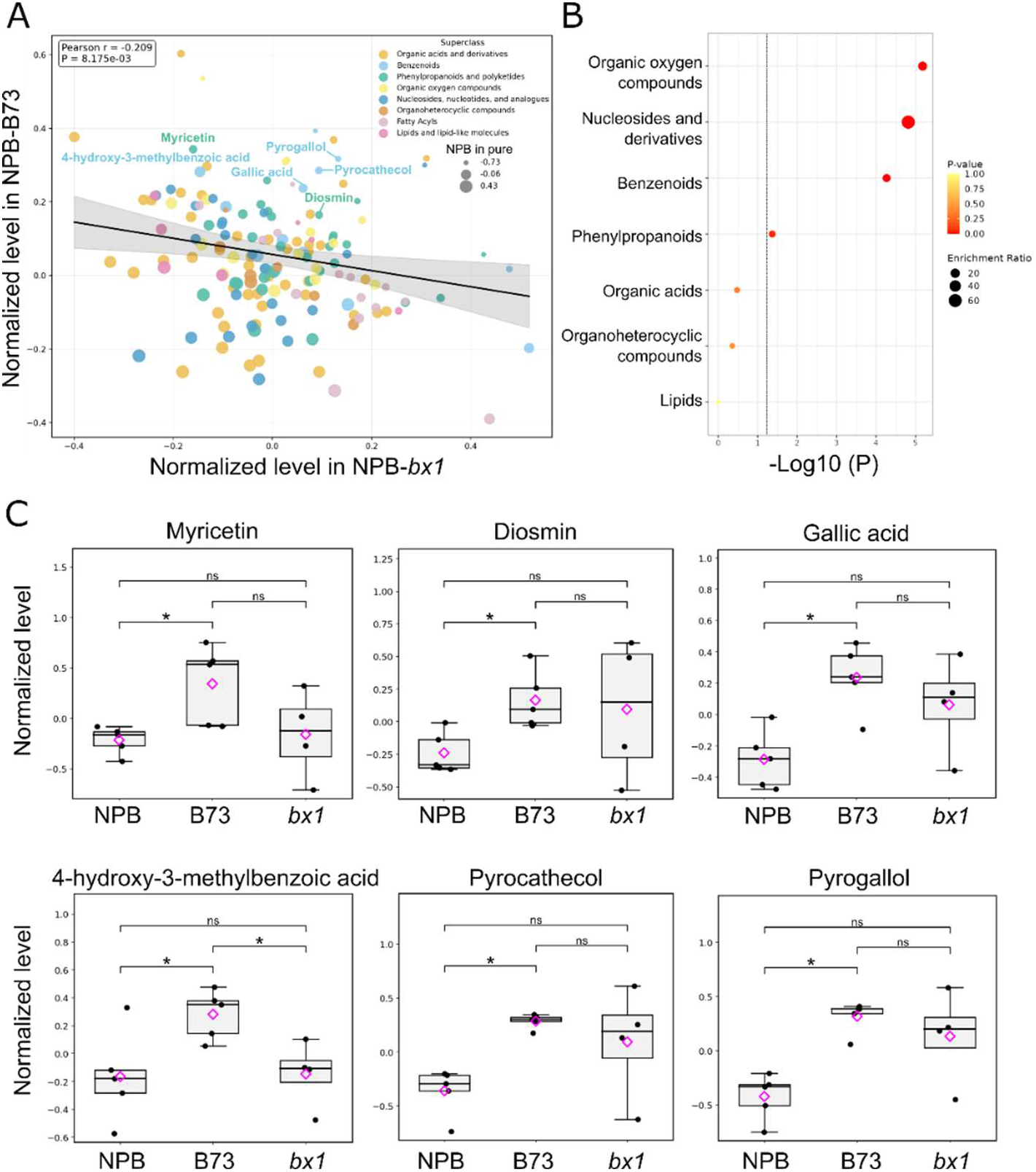
Metabolomic responses of rice leaves to a BX cocktail. **A.** Normalized metabolite intensities in *Oryza sativa* cv. Nipponbare (NPB) leaves co-cultured with wild-type maize (B73) or the benzoxazinoid-deficient mutant *bx1*. The black line represents the linear regression, and the grey shadow indicates the 95% confidence interval. Metabolites detected by HPLC-MS/MS were annotated using an authentic standard library. Each point is colored according to its ClassyFire structural superclass, and dot size reflects metabolite abundance in NPB leaves grown in pure culture. **B.** Metabolite structural ontology enrichment test using ClassyFire structural database, performed on metabolites showing significant changes in NPB when co-cultured with B73 compared with pure and *bx1* mutant conditions. **C.** Normalized intensities of specialized metabolites significantly altered in NPB when co-cultured with B73 relative to pure and *bx1* mutant conditions. Magenta diamond indicates the mean. * P<0.05 according to Mann-Whitney U test followed by Benjamini-Hochberg correction.

### BX-dependent, chromatin-based regulation of polyphenol biosynthesis genes *via* H3K27 acetylation

BXs modulate chromatin activity through inhibition of histone deacetylases in *in vitro* plants (Venturelli *et al*., 2015), likewise, other metabolites structurally linked to BXs in animals (Reddy *et al*., 2004; Sixto-López *et al*., 2020). Hence, we tested whether naturally exuded BXs could function through their reported *in vitro* molecular mechanism. Western blot analysis revealed increased global levels of H3K27 acetylation (H3K27ac) in roots of rice co-cultivated with B73 compared to pure or *bx1*, albeit to a lesser extent than treatments of rice in pure with sodium butyrate, a *bona fide* histone deacetylase inhibitor. In contrast, the level of H3K27ac remained unchanged in leaves (Figure 6A). Chromatin immunoprecipitation using an anti-H3K27ac antibody followed by qPCR showed enhanced recovery of three genes related to phenylpropanoid biosynthesis in roots of rice exposed to BXs, including *OsPAL1*, *OsCHI3* and *OsF3’H* (Figure 6B). This enrichment correlated with higher transcript levels of these genes in rice roots (Figure 6C), suggesting transcriptional activation associated with chromatin activation. In contrast, chromatin acetylation and associated gene expression at phenylpropanoid genes did not significantly change in response to BXs in rice leaves (Figure S1 and S9), consistent with unchanged *OsPAL1* expression in leaves (Figure S1B). The results support a model in which BXs hyperactivate root chromatin at metabolism genes and thus metabolic activity in rice roots, while chemical changes in leaves are due to secondary signaling in response to root changes rather than direct effects of BXs.

**Figure 6.**
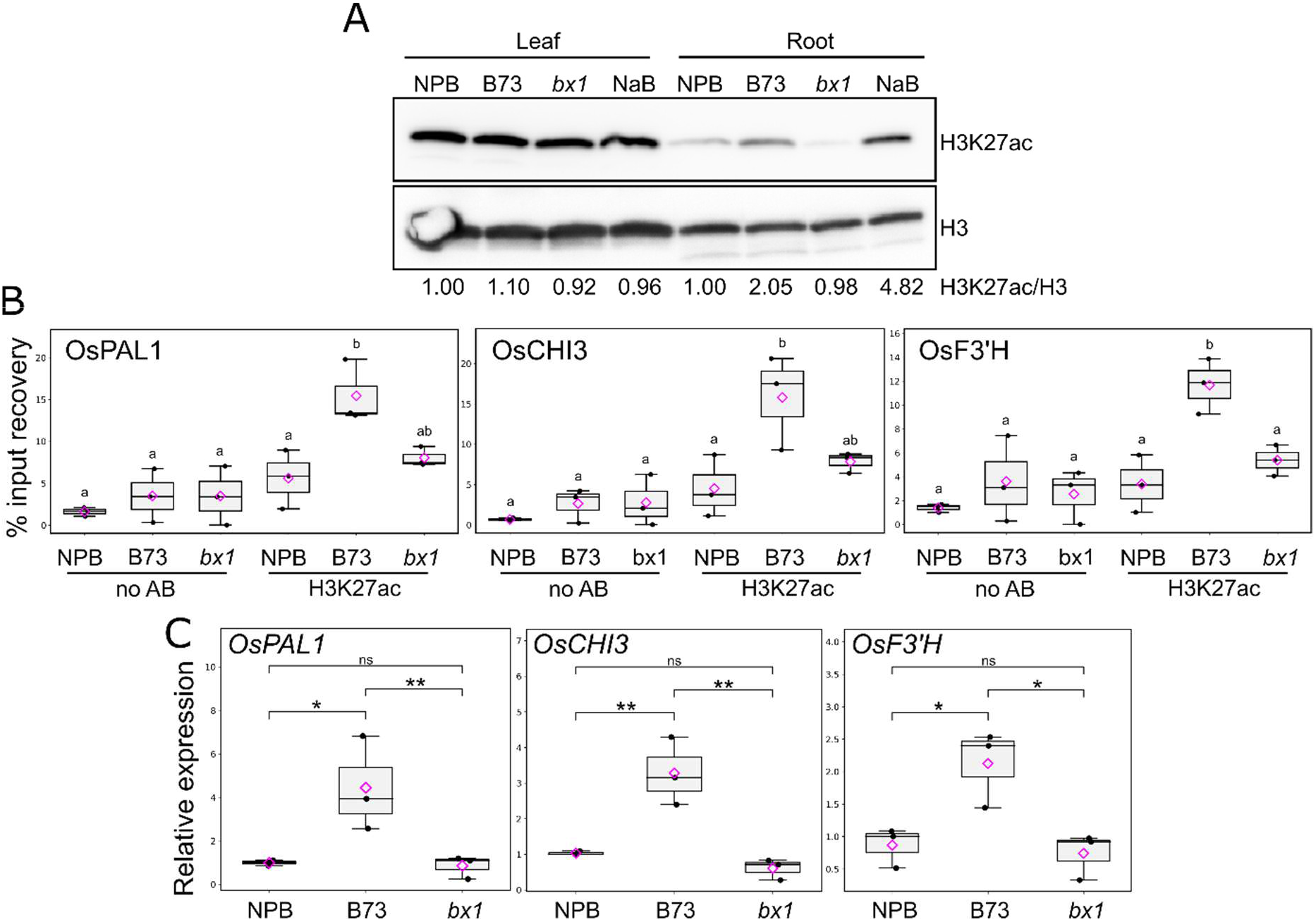
Probing the histone deacetylase inhibitor activity of benzoxazinoids at phenylpropanoid biosynthesis genes. **A.** Western blot with anti-H3 and anti-H3K27ac antibodies to probe chromatin samples of *Oryza sativa* cv. Nipponbare (NPB) in pure or co-cultivated with wild-type maize (B73) or the benzoxazinoid-deficient mutant bx1 at 0h before infection. Pure NPB treated with the histone deacetylase inhibitor sodium butyrate (NaB) was included as a positive control. H3K27ac signals were normalized against H3 within each sample, and pure NPB used as a control in each tissue separately. Representative result of three independent replicates. **B.** Chromatin Immunoprecipitation-qPCR experiment with metabolic gene-specific primers on rice root chromatin in pure or co-cultured with B73 or *bx1*. Samples submitted to blank immunoprecipitation without antibodies (no AB) were included to evaluate background DNA contamination in pull-downs. Different letters indicate statistical differences according to ANOVA and Tukey HSD post-hoc test. **C.** Relative gene expression in rice roots from RT-qPCR assays (ΔΔCt method, normalized against *OsUBQ* and NPB in pure) at 0h before infection. Magenta diamond indicates the mean. * P<0.05, ** P<0.01, *** P<0.001 based on ANOVA followed by Tukey’s HSD post hoc test.

## Discussion

Our results suggest that a BXs cocktail exuded by maize roots and imported into neighboring rice roots decreases rice blast disease through a systemic secondary signal targeted to leaves. We provide evidence that rice roots internalise these maize-derived metabolites, globally inducing increased chromatin and metabolic activities. These changes are characterised, but not limited to, by global activation of primary metabolism and contrasted and specific regulation of defense-related specialized metabolism in roots and leaves (Figure 7). In contrast with the well-documented toxic effects of exogenously applied benzoxazinoids at high concentration, we propose that rice-maize interaction follows a hormesis-like logic, which likely applies to a variety of biochemically-mediated organismic interactions.

**Figure 7.**
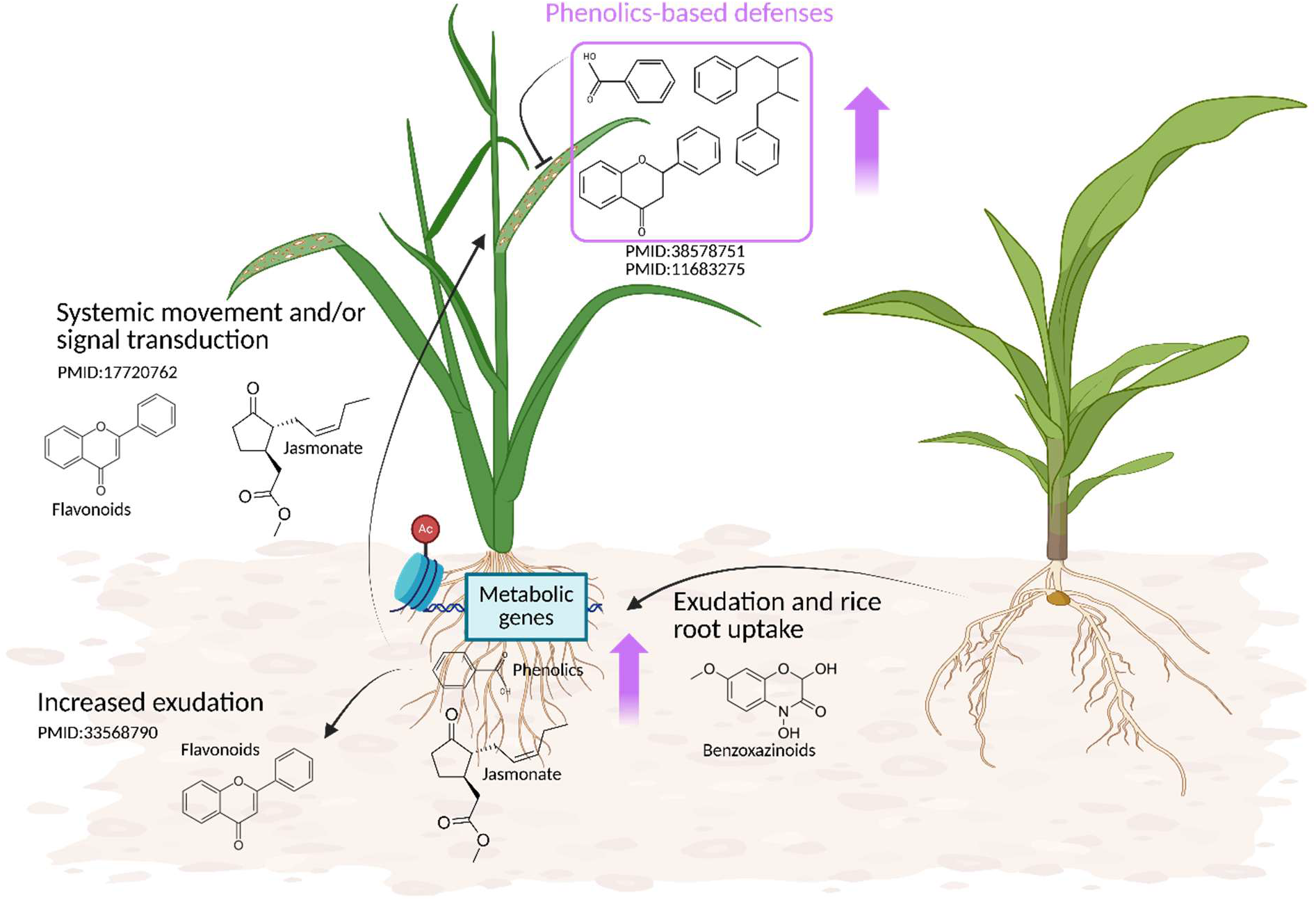
Proposed model of benzoxazinoid-dependent maize-rice interactions leading to enhanced resistance to rice blast. Maize-derived benzoxazinoids are released into the rhizosphere and can be taken up by rice roots, where they modulate chromatin activity, metabolic and defense-related processes. In rice roots, exposure to these compounds is associated with increased jasmonic acid, a known inducer of specialized metabolism. BX internalization leads to chromatin activation at phenolic biosynthesis genes, increased accumulation of simple phenolics and a concurrent decrease of of flavonoids, potentially due to increased exudation and/or systemic redistribution of phenolic compounds within the plant. These root-level responses are linked to the activation of phenolic-based defenses in rice leaves including anti-rice blast compounds such as Myricetin. Therefore, these benzoxazinoid-mediated processes are proposed to induce phenolic defense pathways in rice and thereby attenuate rice blast disease. Created in BioRender. Mathieu, L. (2026) https://BioRender.com/6d8vlb9; agreement number PP29XC06LN.

### Interspecific transfer of maize benzoxazinoids to rice roots might trigger systemic molecular changes

Previous studies have shown that BX-conditioned soils shape microbial communities that in turn affect plant growth and defense responses against a variety of threats (Hu *et al*., 2018a; Schandry & Becker, 2025; Stengele *et al*., 2026). Therefore, BXs are largely considered to influence plant physiology indirectly, by reshaping bacterial and fungal assemblages in the soil (Thoenen *et al*., 2024a). Together with other studies (Venturelli *et al*., 2015; Kong *et al*., 2018; Knoch *et al*., 2022; Gfeller *et al*., 2023), our results support an additional direct mechanism whereby BXs are transferred from maize to rice roots and induce physiological regulations distinct from allelopathic or microbiome-mediated effects.

The mechanisms underlying BX uptake remain unknown (Hama *et al*., 2024), but their accumulation in rice roots suggests uptake from the rhizosphere via passive diffusion or unidentified transport systems. Previous studies detected highly variable amounts of BXs systemically transported from roots to leaves relative to the receiver species analyzed, from a limited quantity of few specific BXs to comparable amounts in roots and leaves of the whole BX cocktail (Macías *et al*., 2014; Hazrati *et al*., 2020; Hama *et al*., 2026; Luo *et al*., 2026). Here, we did not detect any traces of BXs in leaves, albeit detected to high levels in roots (25% of the levels in B73 roots), which argues against a direct toxic effect of the BX cocktail on *M. oryzae* and shows that rice does not systemically transport BXs, or with poor efficiency and hence under our limit of detection. Again contrasting with previous findings where the effects of a BX-structured microbiome were demonstrated, we did not observe convincing changes in systemic acquired resistance gene expression such as *PR* genes, or growth changes in response to BX root uptake in our experimental design (Hu *et al*., 2018b; Stengele *et al*., 2026). However as previously reported in response to BX signaling, we observed increased JA and cathecol amounts (Figure S4A-B) (Kong *et al*., 2018; Richter *et al*., 2024). Together with existing studies, our results support the idea that BX signaling induces receiver species and/or context-specific effects and identifying BX import processes will be an important step toward understanding the extent and selectivity of such interspecific metabolite exchanges in plant communities. Nonetheless as mentioned in the introductory section, we insist that the observations reported here likely depends on the functional composition of the rhizosphere microbiota, as the production of certain BXs depends on specific microbial enzymatic activities (Thoenen *et al*., 2024b).

### Benzoxazinoids trigger metabolic reprogramming associated with chromatin regulation

We show that BX uptake in rice roots triggers metabolome activation, including simple phenolics and core metabolism pathways. This observation correlates with the global increase in H3K27 acetylation, a histone mark associated with active chromatin states and transcriptional activity (Earley *et al*., 2006). Notably, we observed that chromatin hyperacetylation was associated with enhanced transcript abundances of several phenylpropanoid biosynthesis genes, including *Chalcone Isomerase 3*, as reported previously in response to BX exposure in faba bean (Li *et al*., 2016). However, we observed a seemingly opposite effect on cellular levels of the flavonoid class of polyphenols in rice roots that strongly decreased. In fact, previous studies showed that BXs increased flavonoid exudation, which could decrease the cellular pool of flavonoids (Li *et al*., 2016; Dong *et al*., 2022). Together with other studies, our results align with previous observations in various species in response to BXs, where the regulation of flavonoid biosynthesis and/or exudation appears as a hallmark response of BX signaling (Li *et al*., 2016; Leoni *et al*., 2021; Dong *et al*., 2022; Hama *et al*., 2026). Therefore, although BX perception induces species-specific responses to some extent, some physiological responses such as jasmonic acid signaling, simple phenolics based defenses, and increased flavonoid exudation might be conserved. Increased flavonoid exudation in response to BXs might have profound implications given their roles in shaping root-microbes interactions in the rhizosphere (Weston & Mathesius, 2013). Additional work is therefore required to disentangle the interplay between benzoxazinoids- and flavonoids-mediated structuring of the rhizosphere microbiome.

### Root-to-shoot coordination drives the spatial redistribution of phenolic defenses

In contrast with roots, the accumulation of complex phenolic compounds including some flavonoids in leaves, supports the existence of systemic signaling between roots and shoots in response to the BX cocktail. A putative mechanism could involve flavonoids or their derivatives as mobile signals (Buer *et al*., 2007, 2008) that coordinate metabolic responses across tissues in response to BXs. Alternatively, BX perception in roots may activate long-distance signaling pathways involving phytohormones such JA, a potent inducer of specialized metabolism (Figure 7) (47, 48). Consistent with this hypothesis, JA accumulated in rice roots in a BX-dependent manner (Figure S4A), accompanied by a slight but significant BX-dependent increase in *JAZ1* transcript abundance in leaves (Figure S1). From a functional perspective, the spatial regulation of phenolic metabolism may contribute to rice blast suppression because phenolic compounds, including flavonoids such as Myricetin, possess antifungal properties (Zhang *et al*., 2023; Moin *et al*., 2024; Jiang *et al*., 2024). Collectively, our results suggest that a BX cocktail generates root chromatin activation, a root-initiated systemic response leading to the accumulation of defense-related metabolites in distant plant tissues. Our results broaden the recognized roles of BXs as ecological regulators linking plant–plant chemical interactions, chromatin-associated regulation, metabolic reprogramming, and resistance to diseases. More broadly, our results suggest that a hormesis-like logic, whereby compounds historically viewed as toxic can elicit beneficial responses at low or natural exposure levels (Agathokleous *et al*., 2020; Erofeeva, 2022; Kong *et al*., 2024; Mathieu *et al*., 2024; Agathokleous, 2025), may provide a useful framework for understanding molecular plant–plant interactions mediated by root exudates. This perspective opens new avenues for exploiting natural interspecific chemical interactions to develop sustainable crop protection strategies through plant diversification.

## Acknowledgments and funding sources

This work was supported by ANR fundings 20-PCPA-0006 and ANR-24-CE20-7756, and by the SPE INRAE department. We are grateful to Christelle Robert and Matthias Erb for discussions and the sharing of WT and mutant maize seeds. We are grateful to Henri Adreit for seed production. The authors also acknowledge financial support from the Bordeaux Metabolome Facility, MetaboHUB (ANR-11-INBS-0010), and PHENOME (ANR-11-INBS-0012) research infrastructures.

## Author contributions

### Author contributions

1. L. M.: Methodology, Formal analysis, Data curation, Visualization, Writing – original draft, Writing – review & editing; R. P.: Investigation, Methodology, Formal analysis, Data curation, Writing – review & editing; I. B.: Investigation, Methodology, Formal analysis, Writing – review & editing; N. P.: Methodology, Formal analysis, Writing – review & editing; A. R.: Methodology, Metabolomic data acquisition, Formal analysis, Data curation, Writing – review & editing. C. R.: Methodology, Metabolomic data acquisition, Formal analysis, Data curation, Writing – review & editing; P. P.: Methodology, Resources, Supervision of metabolomic analyses, Formal analysis, Writing – review & editing; J.-B. M.: Conceptualization, Resources, Supervision, Funding acquisition, Writing – review & editing; L.-V. M.: Conceptualization, Methodology, Supervision, Project administration, Funding acquisition, Formal analysis, Visualization, Writing – original draft, Writing – review & editing. Laura Mathieu and Rémi Pélissier contributed equally to this work.

