## Supplemental figures for "Interspecific transfer of root-exuded specialized metabolites coincides with chromatin regulation and systemic chemical defense in rice"

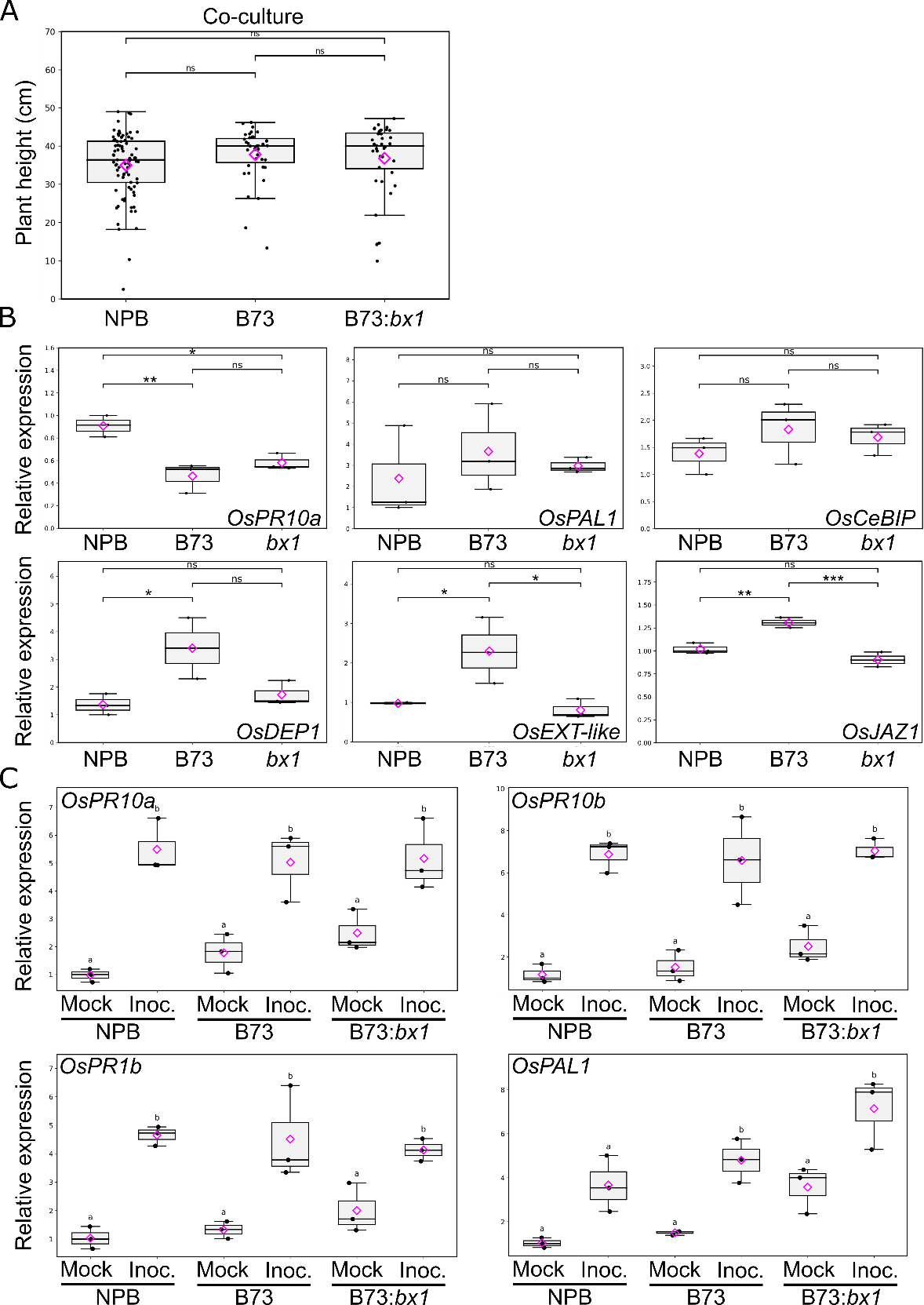

**Figure S1. Growth and defense gene expression of rice in response to pure culture or co-culture with benzoxazinoid-producing maize (B73) or benzoxazinoid-deficient maize (*bx1*).**

**A.** Total plant height of *Oryza sativa* cv. Nipponbare (NPB) measured after three weeks of growth in pure or mixed conditions. ns indicates no significant difference according to pairwise Mann-Whitney U test with Benjamini-Hochberg correction.

**B-C.** Relative gene expression in rice leaves from RT-qPCR assays (ΔΔCt method, normalized against *OsUBQ* and NPB in pure) at **B**. 0h before infection and **C**. 48 hours after mock or *M. oryzae* infection. Magenta diamond indicates the mean. Different letters indicate statistical differences based on ANOVA followed by Tukey HSD post-hoc test.

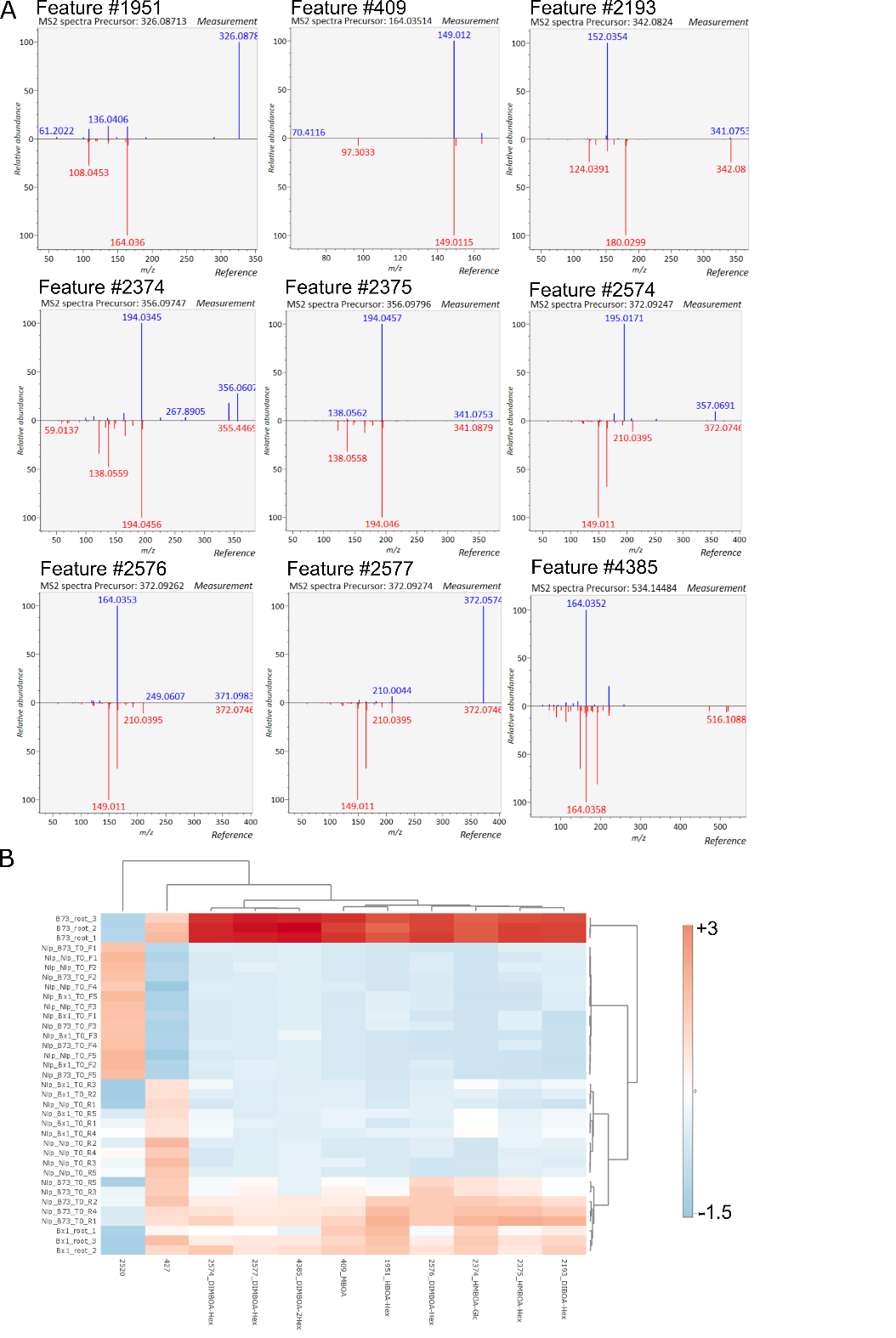

**Figure S2. Identification of benzoxazinoids by tandem mass spectrometry and benzoxazinoid-dependent clustering of rice and maize tissues prior to infection.**

**A.** Fragmentation profiles of detected metabolites (blue) compared with benzoxazinoid standards (red), obtained by tandem mass spectrometry (MS/MS).

**B.** Heatmap showing hierarchical clustering using Euclidean distance and Ward’s linkage for both rows and columns). Rows were centered and scaled to unit variance before visualization) of leaf (red) and root (blue) samples from rice (Nipponbare, Nip) and maize grown either in pure culture (Nip) or in co-culture with the wild-type maize (B73) or the benzoxazinoid-deficient mutant *bx1*.

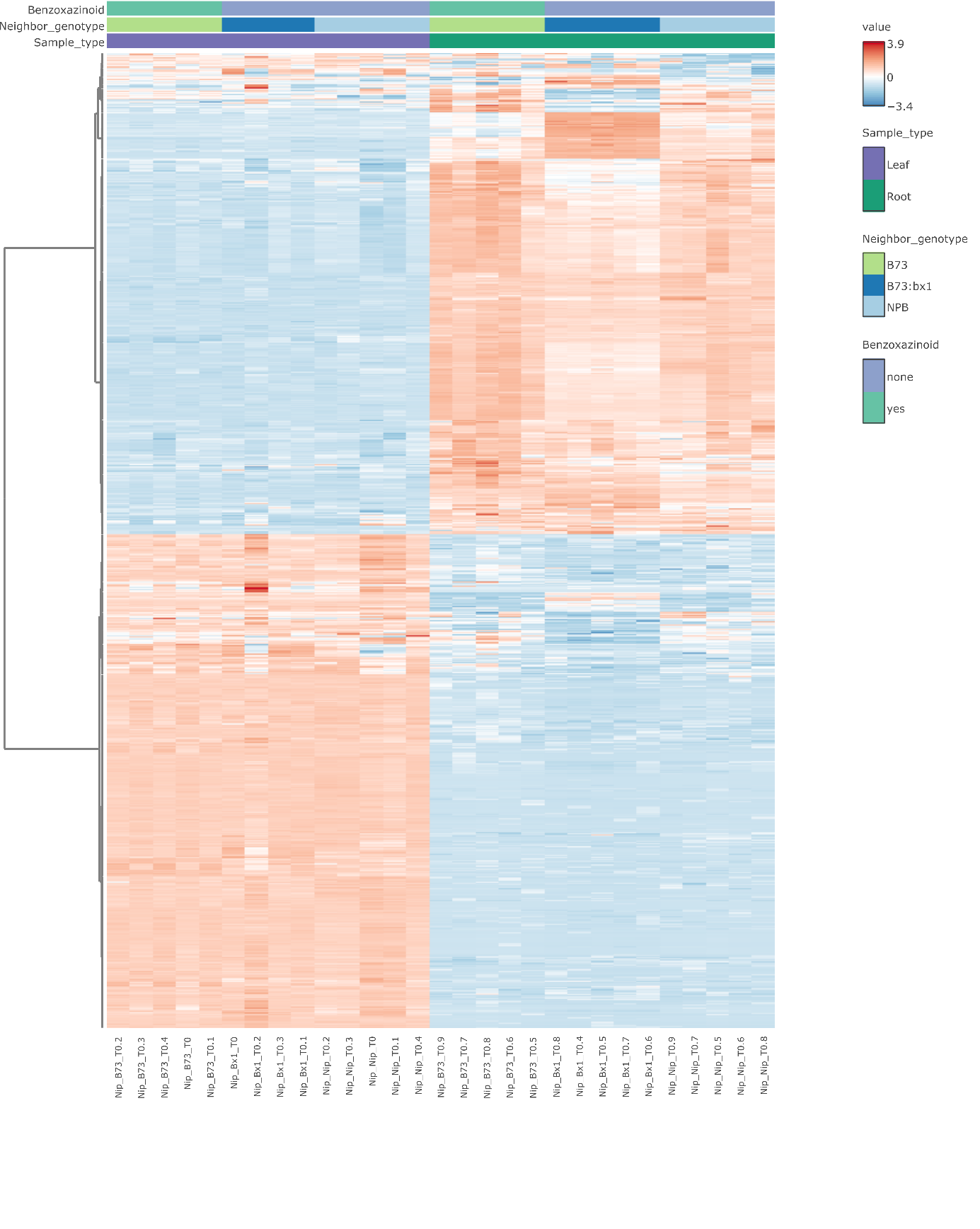
**Figure S3. Hierarchical clustering of rice metabolome datasets.** Metabolite profiles were standardized to feature-wise Z-scores and hierarchically clustered using Euclidean distance and Ward’s linkage. Leaf (purple) and root (green) samples from rice (Nipponbare, Nip / NPB) grown either in pure culture or in co-culture with the benzoxazinoid-producing maize (B73, light green) or the benzoxazinoid-deficient mutant *bx1* (blue).

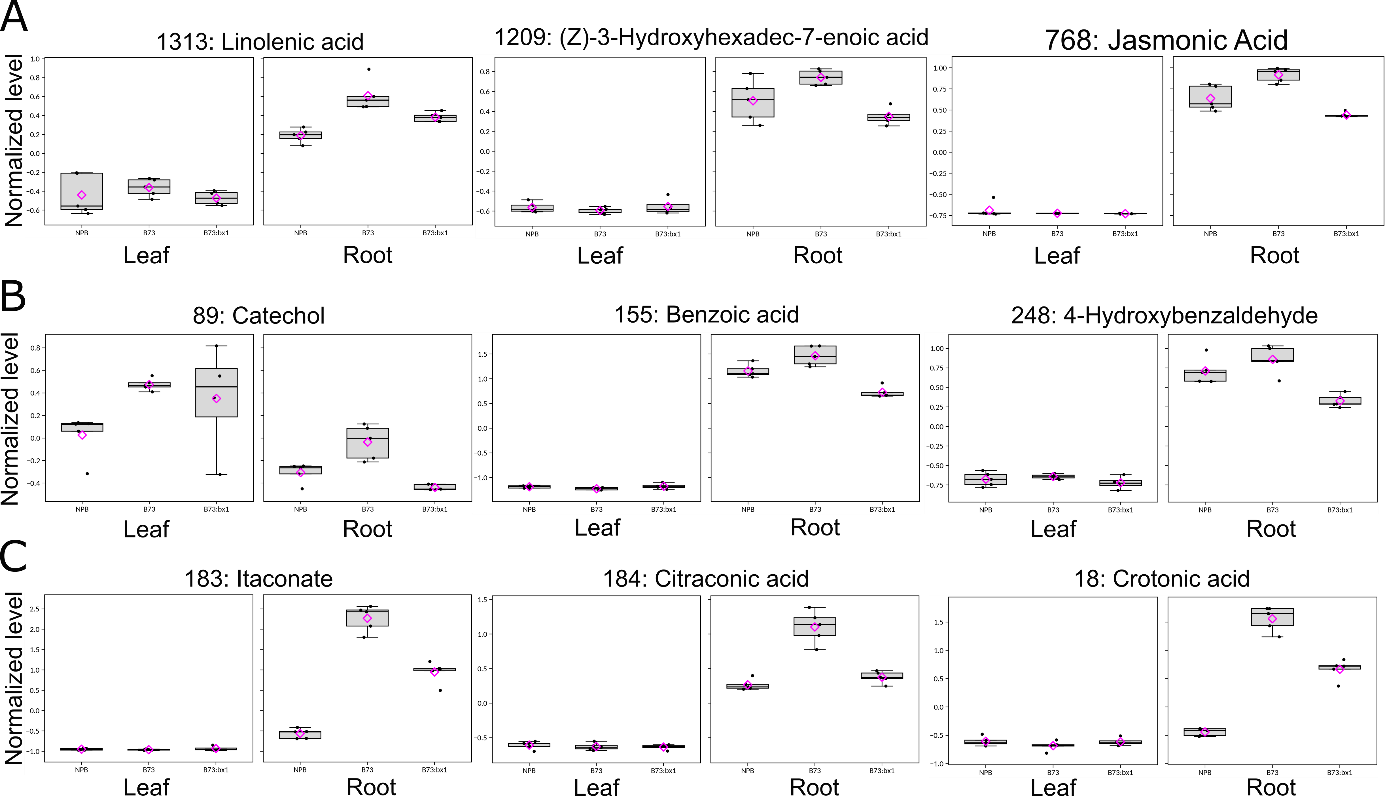

**Figure S4. Representative metabolites identified by generalized linear modelling.** A, Fatty acyls. B, Benzenoids. C, Carboxylic acids

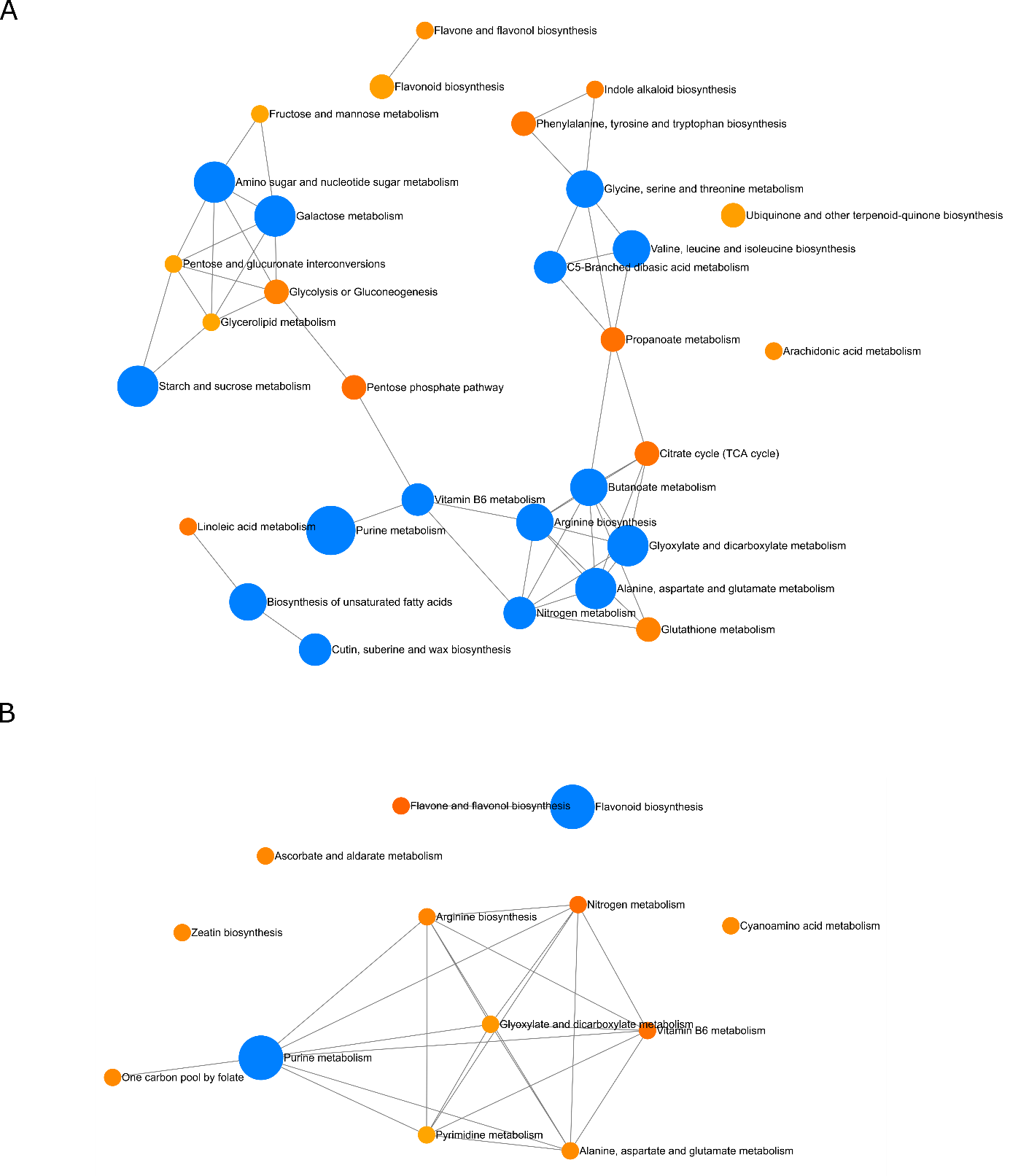
**Figure S5. Metabolic network enrichment analysis in response to benzoxazinoids.** A, Up-regulated and B, down-regulated metabolites were used for pathway enrichment analysis in Metaboanalyst v6.0. Relative-betweenness centrality and hypergeometric test followed by Benjamini-Hochberg multiple testing correction were used for network reconstruction. Significantly enriched pathways are shown as blue nodes with FDR < 0.05.

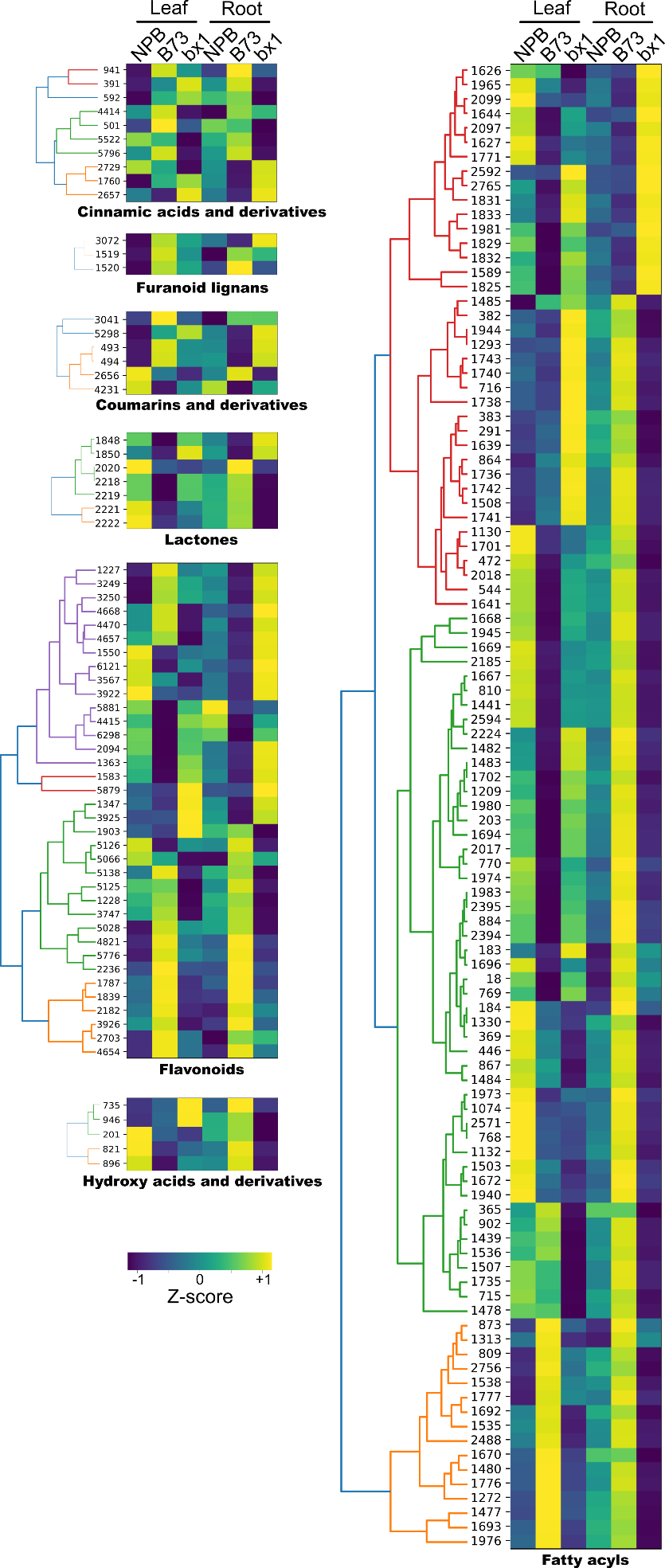
**Figure S6. Heatmap clustering of BX-dependent contrasted metabolic activities in leaves and roots.** Tissue-wise Z-scores were computed for each metabolite and averaged. Clustering was performed using Euclidean distance and average linkage.

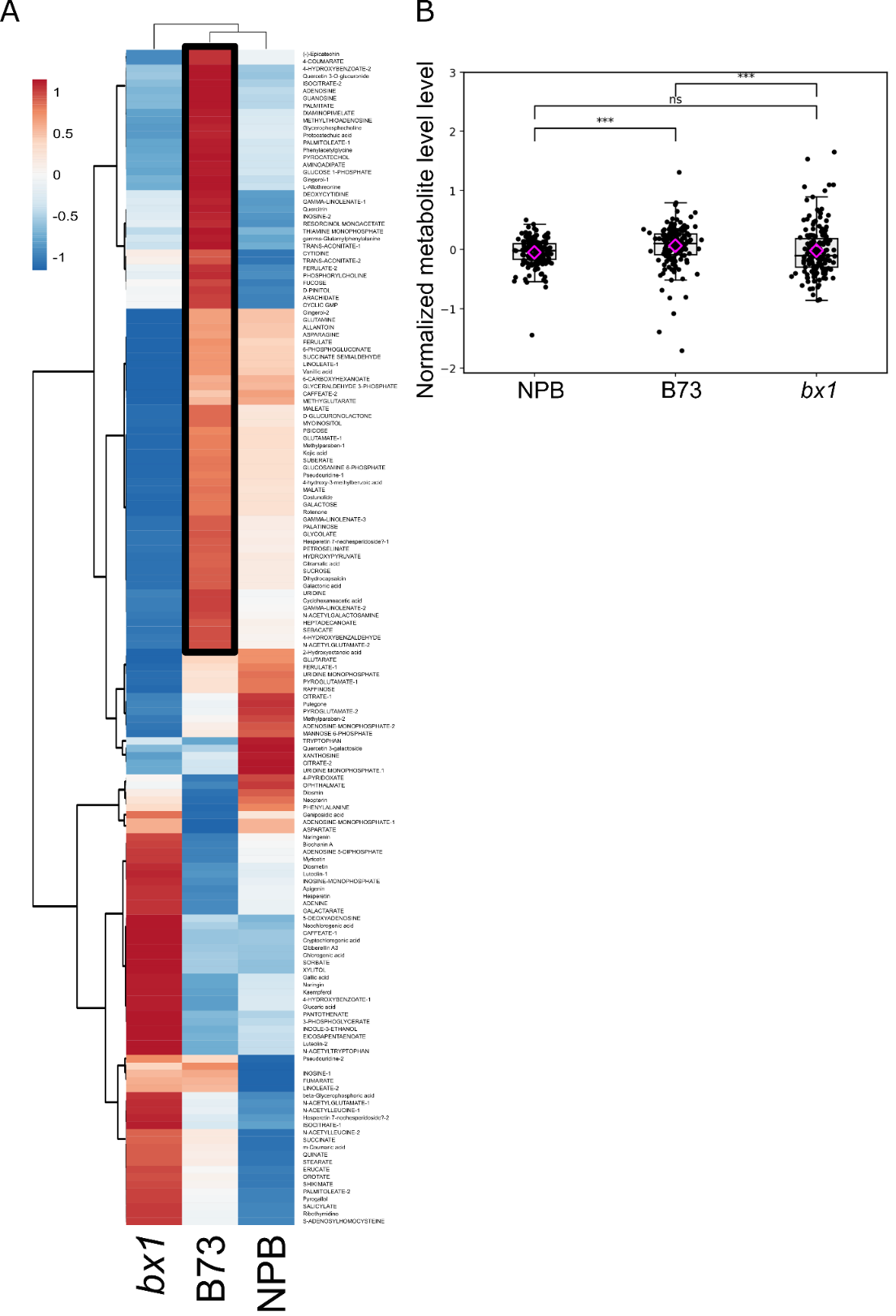

**Figure S7. Global analysis of most confidently annotated metabolites in rice roots in response to BXs.** A, Hierarchical clustering using Pearson correlation distance with average linkage for rows. B, Boxplots represent the mean of normalized levels of all metabolites in roots. Magenta diamond indicates the mean. *** P<0.001 according to pairwise Mann-Whitney U test with Benjamini-Hochberg correction.

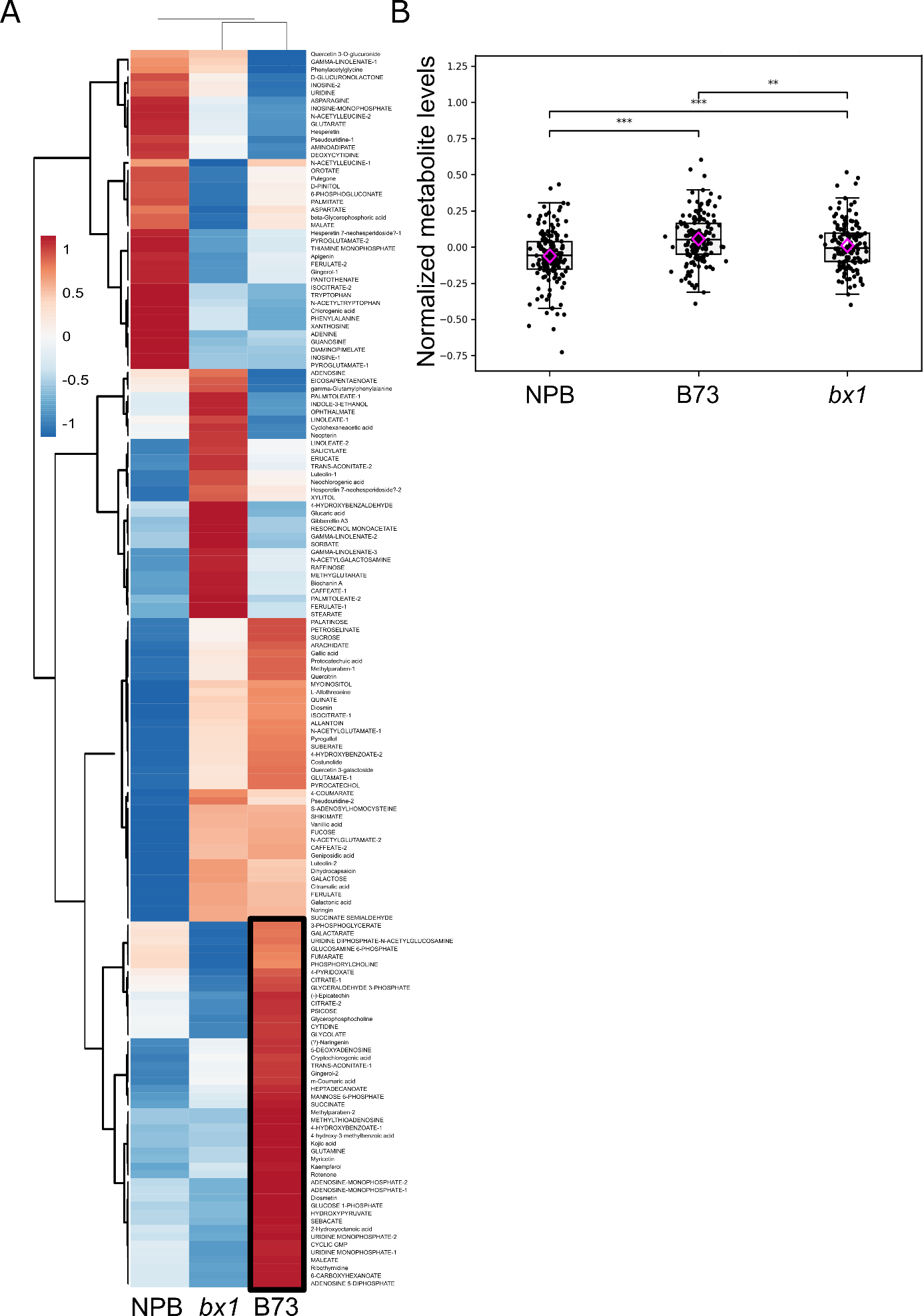

**Figure S8. Global analysis of most confidently annotated metabolites in rice leaves in response to BXs.** A, Hierarchical clustering using Pearson correlation distance with average linkage for rows. B, Boxplots represent the mean of normalized levels of all metabolites in leaves. Magenta diamond indicates the mean. ** P<0.01, *** P<0.001 according to pairwise Mann-Whitney U test with Benjamini-Hochberg correction.

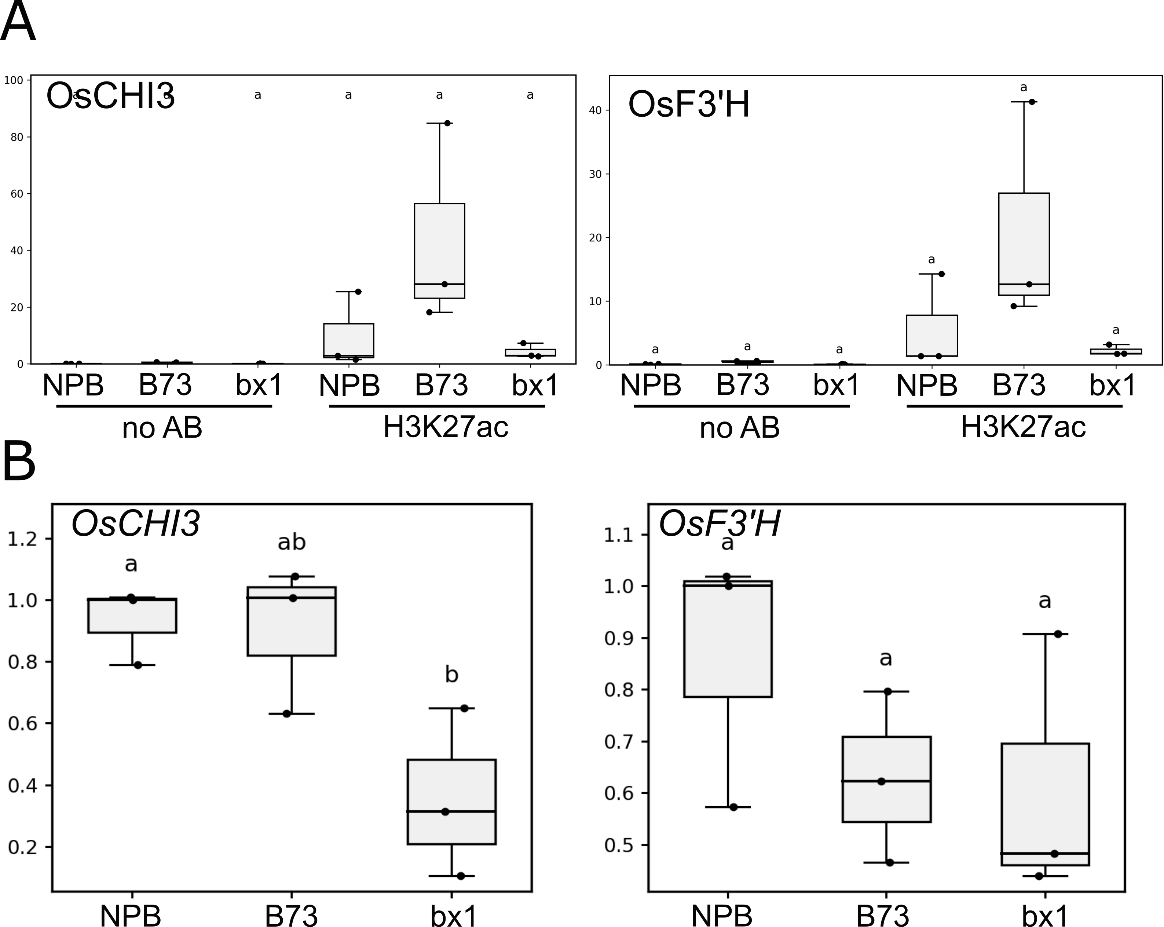

**Figure S9. Expression of polyphenol biosynthesis genes in rice leaves grown in pure or co-cultured with B73 or *bx1*.**

**A.** Chromatin Immunoprecipitation-qPCR experiment with metabolic gene-specific primers on rice leaf chromatin in pure or co-cultured with B73 or bx1. Samples submitted to blank immunoprecipitation without antibodies were included to evaluate background DNA contamination in pull-downs. Different letters indicate statistical differences according to ANOVA and Tukey HSD post-hoc test.

**B.** Relative gene expression in rice leaves from RT-qPCR assays (ΔΔCt method, normalized against OsUBQ and NPB in pure) at 0h before infection. Magenta diamond indicates the mean. Different letters indicate statistical differences based on ANOVA followed by Tukey HSD post-hoc test.
